# Cell-type specific iron content regulation revealed by single-cell iron quantification

**DOI:** 10.1101/2025.06.22.660918

**Authors:** Philip Holdship, Megan R. Teh, Michalina Mazurczyk, Dana Costigan, Giulia Pironaci, Huei-Wen Chuang, Michael B. Zimmerman, Nicole U. Stoffel, Robert G. Hilton, David Price, Jon Wade, Hal Drakesmith

## Abstract

Iron is crucial for cellular metabolism and cell growth. Nevertheless, in humans, both iron deficiency and disorders of iron overload are widespread. How cellular iron content varies depending upon iron availability, and how this influences cell function is poorly characterised. We developed a method to quantify metals in hundreds of cells per minute via single-cell inductively-coupled plasma mass spectrometry (sc-ICP-MS), and used this to explore iron usage by immune cells. Activated murine T-cells exposed to a 625-fold titration of extracellular iron maintained close homeostatic control, with iron content varying by ∼20%. However, these variations strongly correlated with activation characteristics and proliferation. Running sc-ICP-MS downstream of flow cytometric sorting showed that murine T-cells and B-cells *ex vivo* exhibit similar mean and heterogeneity of cellular iron while splenic macrophages contain twice as much iron and more heterogeneous iron content. Finally, activated human B-cells contain ∼10-fold more iron per cell than murine B-cells. We suggest that mechanisms of iron homeostasis impart particular ranges or set-points of iron content to different cell types and activation states, and that small changes in iron content have large effects on cell behaviour. Our methodological advance and consequent findings suggest new approaches to studying the biology of metals.

## Introduction

Iron is crucial for multiple cellular biochemical activities(*1*). The failure to maintain iron homeostasis has significant consequences for human health(*2*). Iron deficiency is the world’s most common micronutrient deficiency, affecting over a billion people worldwide (*3*). Iron deficiency is particularly common and severe in many low-middle income countries(*4*). Iron deficiency can cause anaemia, inhibit growth and development, and impairs immunity(*5*). Iron is crucial for T-cells and B-cells, to support cellular proliferation(*6*, *7*), activation(*8*), and metabolic reprogramming(*9*).

Iron deficiency is characterised by plasma parameters such as ferritin or transferrin saturation, which are proxy assessments of iron stores and iron availability for cells, respectively. However, the manifestations of iron deficiency depend on the iron content of cells and how cellular functions are altered by lack of iron. Our knowledge of how iron is distributed between different cell types and how cells respond to different levels of iron availability is currently limited(*10*). This primarily reflects the inherent analytical difficulties in accurately measuring small variations in cellular iron levels amongst representative populations of cells.

Currently, cellular iron contents are typically determined by bulk dissolution, followed by analysis of the resultant solutes by various atomic spectroscopy techniques (for example, inductively coupled plasma atomic emission spectrometry(*11*)). Bulk approaches cannot assess heterogeneity among and between cell populations. This may lead to uncertainties in the derived mass per cell values as well as lack of understanding of how iron is partitioned across cell populations. Analyses of individual cells by nanoSIMS and synchrotron µXRF, enables accurate elemental quantification on a per-cell basis. However, these microanalytical approaches are limited by the number of cells that may be analysed, the fixing procedures required and, in the case of synchrotron-based instruments, timely access to the instrument.

An alternative approach is single-cell inductively coupled plasma mass spectrometry (sc-ICP-MS). A suspension of cells is aspirated into an ICP-MS, dispersed sufficiently to be ionised, and subsequently analysed, sequentially. Analytical time intervals are set sufficiently short that cells transiting through the instrument can be discriminated from the background and ion counts arising are integrated to yield a total cellular mass for the element of interest. This allows for the analysis of individual cells at a frequency of hundreds to thousands of cells per minute, sufficient for exploring cell population heterogeneity and providing a potentially rapid screening tool. However, the measurement of iron by this analytical method has been described in only a few research studies using either a tumour cell line (*12*), yeast, alga and red blood cells (*13*), or bacteria(*14*, *15*).

In this study, we optimised sc-ICP-MS for cellular material and cross-validated data against bulk ICP-MS and CyTOF. We assessed single-cell metallomics in populations of several different types of leukocyte (Jurkat cells, murine T-cells, B-cells and macrophages, and human primary B-cells). We also cultured murine T-cells across a titration of iron availability and related subsequent iron content to characteristics of cell activity.

## Results

### Single cell metallomics by sc-ICP-MS validated by titrated intercalation experiments

Rhodium and iridium, in their 3+ cationic forms, are effective metallointercalators when complexed with polycyclic aromatic ligands(*16*, *17*). When conjugated to antibodies, they are employed for cell analyses by CyTOF mass cytometry (*18*). CyTOF allows for multiplexed analyses of single cell populations through measurements of tagged lanthanides/heavy metals (*19*). However CyTOF cannot constrain the divergence of low mass elements out of the highly energetic ion beam by charge repulsion (known as space charge effects) (*20*, *21*), so is currently unable to measure endogenous biometals, such as iron. We used rhodium and iridium intercalated cells to test sc-ICP-MS as a metrological metallomic analytical technique, via comparison with CyTOF.

Titrated concentrations of rhodium and iridium intercalators were added to Jurkat cells, a T-cell line, and uptake was assessed through attogram/cell (10^-18^g) mass range by sc-ICP-MS, CyTOF and bulk ICP-MS (see Fig. 1A). For sc-ICP-MS, the peak area from each cell event was used to determine the mass of metal per cell, where we harnessed high-frequency scanning to measure metals as mass-spectral fingerprints with minimal detector exposure times (dwell times), at intervals that were much lower than cell event durations (up to frequencies of 100,000Hz) (see Table 1). This enabled us to rapidly collect highly precise single-cell metallomics from whole cell populations, where hundreds of cells per minute were measured from the highest four titrants from each experiment (Table S1).

**Fig. 1.**
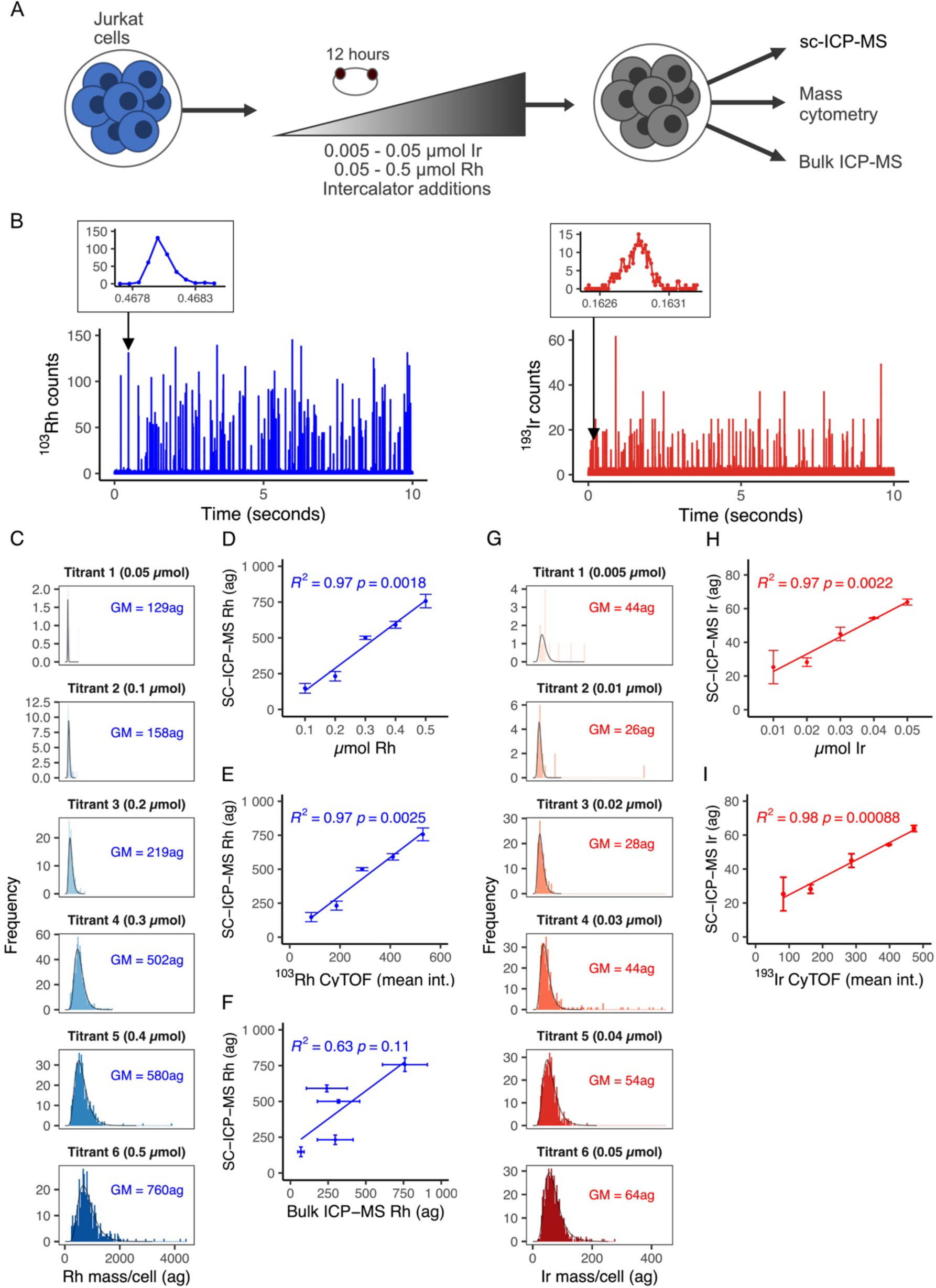
Validation of sc-ICP-MS from rhodium (blue) and iridium (red) intercalation experiments. **(A).** Experimental design for validation of sc-ICP-MS by a metal intercalation approach; **(B).** Excerpts of time-resolved mass spectra of intercalated Jurkat cells, intercalated with 0.3µmol rhodium (left) and 0.04µmol iridium (right). Each spike represents a metallomic event, where their individual areas correlate to mass of rhodium or iridium within each measured cell. The metallomic plumes from each metal take ∼0.0004 – 0.0005s to transit through the instrument (shown in the inset peaks). The inset peaks also illustrate the resolutions achieved for the event profiles, where measurements were taken at intervals of 75µseconds (rhodium) and 10µseconds (iridium); **(C).** Mass-frequency histograms presenting single cell metallomic distributions from the rhodium intercalation series. Mass quantified in attograms (ag) (10^-18^g). Geometric means (GM) of each population are annotated per facet; **(D-F).** Correlation plots of sc-ICP-MS (geometric means from rhodium experiment) versus intercalation molarity, mean CyTOF intensities and bulk ICP-MS analysis, respectively; **(G).** Mass-frequency histograms presenting single cell metallomic distributions from the iridium intercalation series (GM = population geometric means); **(H-I).** Correlation plots of sc-ICP-MS (geometric means from the iridium experiment) versus intercalation molarity and mean CyTOF intensities, respectively. Error bars shown in correlation plots present 2-sigma precision for all sc-ICP-MS data points [n=4]. Error bars from bulk ICP-MS comprise a combination of ICP-MS and cell counting uncertainties, combined using the product rule.

Fig. 1B shows metallomic data taken from the rhodium and iridium sc-ICP-MS analyses, where each individual spike represents a single cell event. The peak insets demonstrate a cell event, where the beginning, end and apex of the example event are clearly defined (Fig. 1B insets). Low detector dwell times have been suggested to adversely affect sc-ICP-MS data quality(*22*), primarily from compromised signal intensities and an increased risk of only partially measuring cell events. However, we demonstrate that the detailed peak profiles, in addition to extremely low background baselines, instead provided enhanced analytical figures of merit (transferable metrics for analytical performance (*23*)), such as linearity and precision. Moreover, high-frequency scanning also facilitates filtering out fragmented/ partly detected cells versus real cell events, where the former transit through the instrument with distinguishably shorter event times than intact cells (Fig. S1).

Mass frequency histograms from rhodium and iridium sc-ICP-MS measurements of each intercalated condition (Figs. 1C, G) are shown with optimised bin sizes and lognormal fittings. Background corrections were also applied (supplementary materials). Successive increases in metal mass per cell were found with increasing intercalation concentration in 10^-18^g mass ranges. The mass range on the x-axis is extended >2 times beyond the fitted distributions, to show the negligible occurrence of outliers. This confirms lack of detections of doublets (cell events that include the concurrent measurement of two cells within the same measurement window), which if detected would significantly impact accuracy of metallomic distributions. Heterogeneity of cellular metal uptake was well-defined, where interquartile ranges of 48ag (0.1µmol) to 385ag (0.5 µmol) for rhodium; and 7.5ag (0.01 µmol) to 36ag (0.05 µmol) for iridium (Table S1). We plotted population geometric means of metal/cell against the titrated intercalator molarity (Figs. 1D, H), showing close correlations (R^2^ >0.97) and p-values (0.002).

Duplicate and triplicate aliquots were measured by CyTOF and ICP-MS, respectively, to compare signal outputs. Excellent agreements were found between geometric means of metallomic distributions gained from the sc-ICP-MS and arithmetic means from CyTOF, for both rhodium and iridium experiments (Figs. 1E, I; R^2^ >0.96 and p-values ≤0.002). A similar correlation plot comparing sc-ICP-MS to bulk ICP-MS from the rhodium experiment revealed a weaker correlation (Fig. 1F;l R^2^ = 0.63, p-value = 0.11). The contrast between the single cell and bulk analytical techniques is driven by the considerably enhanced detection capability from the former method. Notably, bulk ICP-MS performed poorly when analysing lower-level iridium titration, reporting iridium content below the limit of detection.

This data demonstrates that sc-ICP-MS accurately and quantifies cellular metal content in the attogram range for individual cells, measuring hundreds of cells per minute. We then used sc-ICP-MS measure biologically relevant metals.

### Iron analysis by sc-ICP-MS requires chemical resolution for accurate detection

We assayed cellular iron and calcium in CD8+ T-cells taken from murine lymphoid tissues. Measurement of iron and calcium by ICP-MS is complicated by other molecular/atomic entities that can share the same mass, for example ^40^Ar^16^O^+^ and ^40^Ca^16^O^+^ for ^56^Fe^+^, and ^40^Ar^+^ for ^40^Ca^+^. To circumvent this, after the samples were introduced into the ICP-MS and the distinct ions generated, ammonia (NH_3_) gas was used to inhibit such interferences through chemical reduction. This method is effective for selective neutralisation of these interferences, when low quantities of NH_3_ are used; in excess NH_3_ can cause considerable energy losses to the analyte ions (*24*). This is particularly important for sc-ICP-MS, as NH_3_-intensive approaches can induce significant time-elongations to plume events –increasing probability of recording doublets (*12*). In contrast, our charge-transfer method with lower NH_3_ flow rates mitigated this effect, where only minimal peak extensions were found (∼1ms peak width for iron compared to 0.5ms for rhodium where no gas was used) (compare Figs. 2A and 1B, respectively). This is 6-fold less elongation than that reported by a mass-shift approach (*25*), so that the probability of detecting doublets remains negligible.

**Fig. 2.**
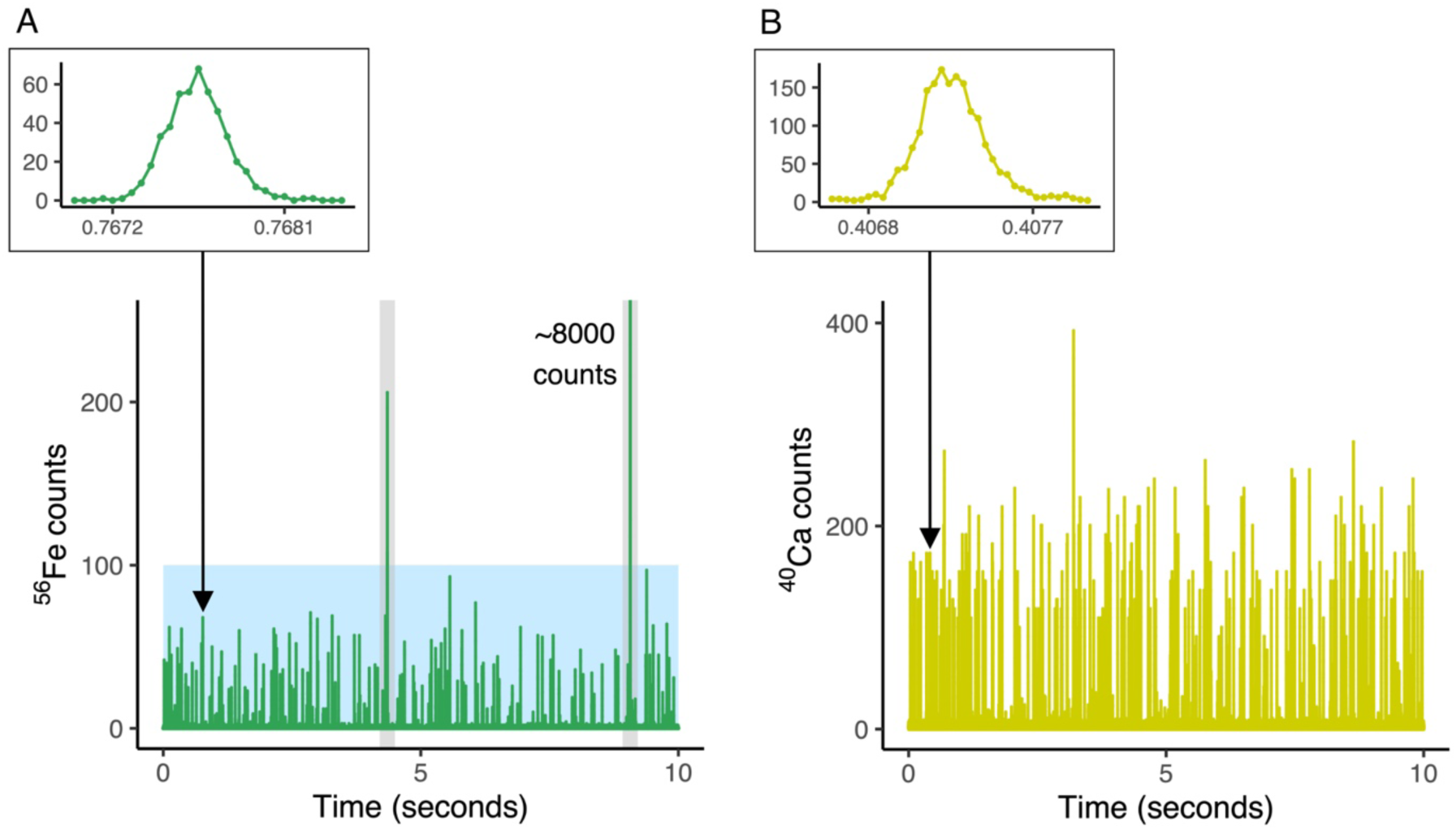
Time-resolved single cell mass spectra measured by sc-ICP-MS from murine T-cells. **(A).** Time-resolved mass spectra from Fe sc-ICP-MS analysis of murine CD8+ T-cells; **(B).** Time-resolved mass spectra from Ca sc-ICP-MS analysis of murine CD8+ T-cells. Each spike in both **(A)** and **(B)** represents an individual metallomic event, where their individual areas correlate to mass of Fe or Ca within each measured cell. The metallomic plumes from each metallomic event take ∼0.001s to transit through the instrument (shown in the inset peaks in both **(A)** and **(B)**). The inset peaks also illustrate the resolutions achieved for the event profiles, where measurements were taken at intervals of 50µseconds (Fe) and 40µseconds (Ca). Note two peaks of very high Fe counts (highlighted with grey bars) – see text. All other peaks are within a similar range (<100 counts) – constituting cell events (highlighted in blue). All measurements conducted using the PerkinElmer NexION 5000 ICP-MS.

A further concern was the appearance of sporadic high-mass events in the iron sc-ICP-MS data (eg peaks within grey shaded bars in Fig. 2A – peaks >200 counts at ∼4 seconds and ∼9 seconds). These intermittent peaks exhibit similar event-widths to cell events but are not seen in the calcium data. We compared the frequency of recorded events (iron vs calcium) – from both unfiltered (all data) and filtered (inclusive of those only within the lognormal distributions, excluding high-mass events) analyses. The number of events recorded from the filtered iron data strongly correlate with similarly filtered calcium results (*R^2^* = 0.81, see Fig. S2); contrasting with a weaker correlation obtained from the amassed iron (cells + high-mass event and calcium data (*R^2^* = 0.57, see Fig. S2). This indicates the high mass iron events were not cells but likely iron oxide precipitates and were subsequently excluded from analysis; calcium high mass events were likely absent, as calcium impurities are soluble at neutral pH.

The calcium data from the cells was additionally used to determine the transport efficiency of our sc-ICP-MS analysis (see Fig. 2B) – the proportion of cells entering the plasma and being detected by the ICP-MS over the entire cell concentration in the measured aliquot, by the particle frequency method (*26*). Signal intensities captured from the apex of peaks were over 30 times higher than the average baseline readings adjacent to the peak [n=17] (see inset to Fig. 2B). From these data, we determined a mean calcium-derived transport efficiency value of 12%, and for iron (after filtering for cell events) the value was 13%. With these methodological protocols in place, we assessed cellular iron content *in vitro* and *ex vivo*.

### Relationship of extracellular iron availability to cellular iron content

CD8+ T-cells were derived from three mice and activated for 48 hours *in vitro* in iron-free media supplemented with titrated concentrations of holotransferrin (transferrin protein with two iron atoms bound) from 0.001mg/mL to 0.625 mg/mL (Fig. 3A). T-cells exposed to these different iron conditions exhibit profound changes in metabolism, mitochondrial activity and proliferation (Teh et al, *Nat Comms* Accepted and https://www.biorxiv.org/content/10.1101/2024.02.01.578381v1.abstract). Analysis of the cells by sc-ICP-MS generated mass frequency histograms of background-corrected single-cell iron data (Fig. 3B), presented with optimised bin sizes, lognormal fittings and annotated geometric mean values. Similar to the intercalation experiments, minimal outliers were found outside the population distributions, again showing negligible occurrence of cell doublets.

**Fig. 3.**
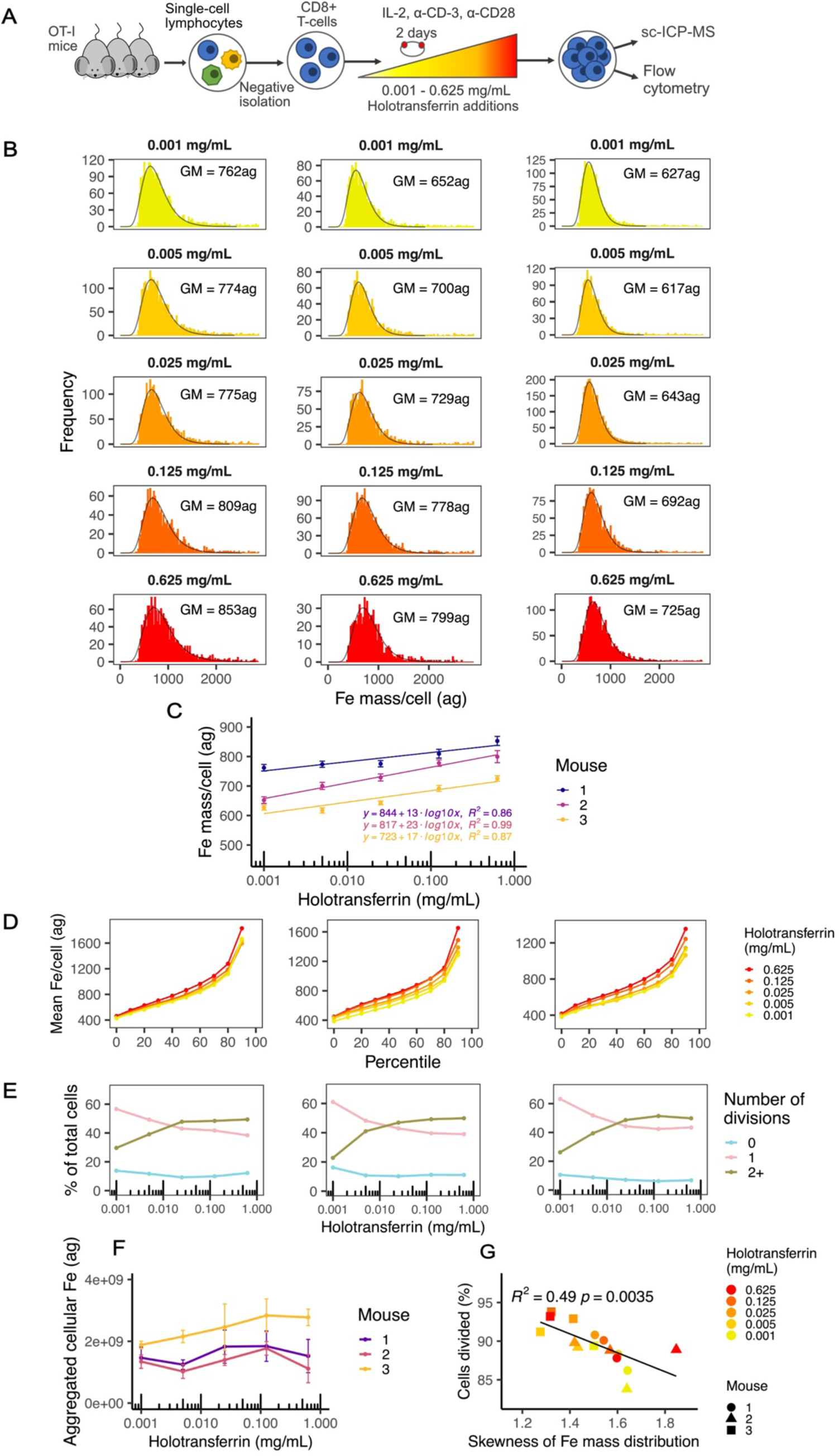
Iron content of murine T-cells exposed to different iron concentrations. **(A).** Experimental design for the assessment of iron in murine T-cells that were exposed to iron media conditions ranging from 0.001mg/mL – 0.625mg/mL holotransferrin; **(B).** Histograms showing the mass-frequency distributions of iron mass/cell for each holotransferrin condition from cells taken from each of three mice. Mass distributions are shown in attograms (ag) (10^-18^g). Geometric means (GM) of each population are annotated per facet; **(C).** Correlation plot presenting the geometric means from each distribution versus the iron condition for each mouse (with a log_10_ x-axis). Logarithmic trend lines, associated equations and *R*^2^ values are displayed for each profile. Error bars show the 95^th^ confidence intervals; **(D).** Line plots showing the mean iron mass/cell per tenth percentile for each holotransferrin condition from each mouse; **(E).** Line plots presenting proportions of cell proliferation activity per iron condition (percentage of total cells dividing 0, 1 or 2+ times) with log_10_ x-axes from each mouse; **(F).** Line plot presenting the mean total amount of iron consumed by the cells from the three mice for each iron condition (with a log_10_ x-axis). Error bars show the range of total iron calculated per mouse; **(G).** Scatter plot presenting the inverse correlation between the magnitude of cell proliferation during the live culturing period against the skewness of the single cell metallomic distributions for iron.

These extrapolated geometric mean iron contents per cell (for each condition and each mouse), were plotted against media iron concentrations (Fig. 3C). Notably, geometric mean iron content per cell increased ∼20% over a 625-fold difference in holotransferrin, demonstrating strong maintenance of cellular iron homeostasis in the face of a range of extracellular iron availability. Nevertheless, over this 20% increase there was a close correlation between increase in iron content and extracellular iron availability (Fig 3C).

Heterogeneity of cellular iron content within each population was evaluated from interquartile ranges (see Fig. S3 and Table S3), indicating increased heterogeneity with exposure to increased iron concentrations. Each population was also segregated into percentiles, and means of each 10^th^ percentile were calculated and plotted per mouse in Fig. 3D. Between the 10^th^ and 70^th^ percentiles smooth trends reflect tight lognormal distributions in iron per cell (see Fig. 3D), which increase in accordance to their iron condition. However, beyond the 70^th^ percentiles, significant escalations in the mean cellular iron occur in every condition, showing the top 20% of cells within each condition contain discrete elevations in iron levels over the rest of the population. Furthermore, the rapid increases in average iron/cell within this range do not occur linearly, where distinctly higher elevations were found for the higher conditions (particularly 0.625mg/mL).

During the 2-day culturing period for this murine T-cell experiment, live cells continually acquired iron and proliferated (Table S4). To examine this effect, we analysed parallel aliquots of cell cultures by Flow Cytometry to assess cell division. The proportions of cells within each population that did not divide, divided once, or divided two or more times over the 48-hour period, are presented in Fig. 3E. The results show increased proliferative activity in the higher iron-bearing conditions, particularly the proportion of cells undergoing 2+ divisions. Using live cell count data and geometric means of iron content/cell, we quantified the total amount of iron used by the cell populations (see Fig. 3F). Interestingly, the peak iron yield associated with the penultimate iron concentration (0.125mg/mL) – it is possible that the highest iron concentration contained some non-transferrin iron that affected cell viability. Across the culture conditions from this experiment, the skewness of the single cell iron mass distributions (proportional to heterogeneity) inversely correlated with the proportion of cells that had divided (Fig. 3G). This elevated skewness is likely due to an increase in the relative proportion of cells that accumulate higher levels of iron within a population that has divided less as a whole.

### Single cell iron metallomic data correlates with markers of cell state

In parallel with sc-ICP-MS on murine CD8+ T-cells, we assayed cell surface CD71 (transferrin receptor) and CD25 (IL-2 receptor) proteins. CD71 mediates transferrin-bound iron into eukaryotic cells. CD71 is highly expressed by activated T-cells, but its synthesis is also regulated by intracellular iron content downstream of intracellular iron-sensing mechanisms (*27*). Relatively iron deficient cells have more surface CD71 to better capture any available extracellular iron(*9*). CD25 is the alpha chain component of the IL-2 surface receptor on T-cells. IL-2 signalling via CD25 promotes T-cell growth and facilitates their differentiation after activation(*28*).

Using flow cytometry, we measured CD71 and CD25 by assessing mean fluorescence intensity (MFI) of anti-CD71 or anti-CD25 fluorophore-labelled antibodies. MFI was plotted against geometric means of iron mass from each cell distribution measured by sc-ICP-MS in parallel aliquots (Figs. 4A, B). We observed an inverse correlation between iron content per cell and CD71 MFI (Fig 4A). Therefore, the small differences in mean cellular iron detected by sc-ICP-MS between conditions reflect biologically meaningful changes that are sensed within cells and regulate synthesis of CD71. CD25 expression positively correlated with iron content (Fig. 4B), in line with the importance of iron acquisition for cellular activation and growth. We plotted CD71 and CD25 versus the geometric means of calcium mass from each cell distribution measured by sc-ICP-MS (see Figs. 4C, D). There were no correlations of calcium content with CD71 or CD25, showing the relative specificity of the relationships between iron content and T-cell activation.

**Fig. 4.**
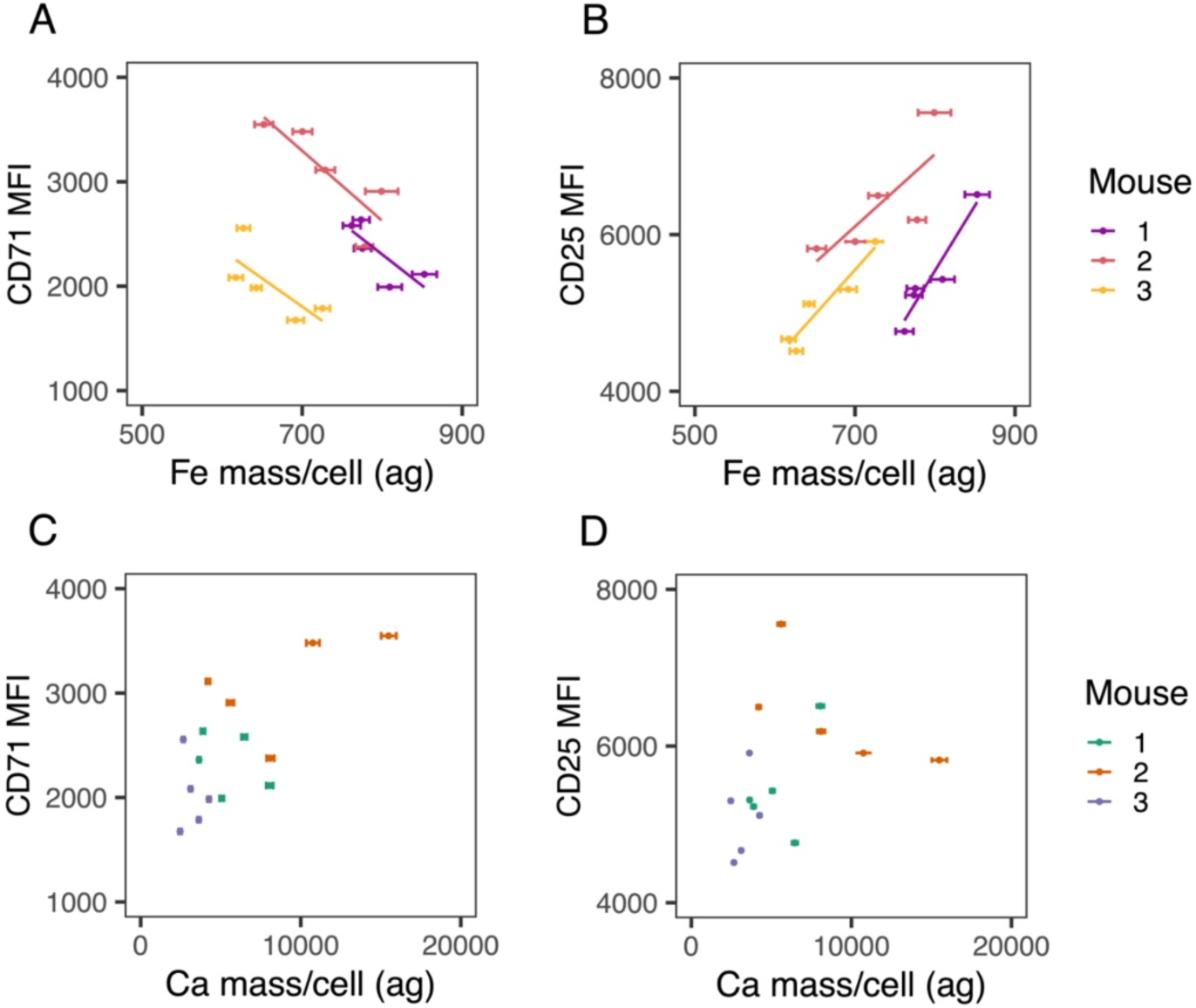
Correlation of surface protein marker expression determined by Flow Cytometry against sc-ICP-MS derived single cell metallomic contents. **(A).** Mean fluorescence intensity (MFI) of CD71 (transferrin receptor) versus the geometric mean single cell Fe content (ag) for each population distribution, from each holotransferrin condition; **(B).** Mean fluorescence intensity (MFI) of CD25 (alpha chain component of the IL-2 surface receptor on T-cells) versus the geometric mean single cell Fe content (ag) for each population distribution, from each holotransferrin condition; **(C).** Mean fluorescence intensity (MFI) of CD71 versus the geometric mean single cell Ca content (ag) for each population distribution, from each holotransferrin condition; **(D).** Mean fluorescence intensity (MFI) of CD25 versus the geometric mean single cell Ca content (ag) for each population distribution, from each holotransferrin condition. Error bars present the 95^th^ confidence intervals for each metallomic distribution determined by sc-ICP-MS.

### Direct *ex vivo* measurement of cellular iron content and heterogeneity

To measure iron content of cells in a more physiological environment, bulk lymphocytes derived from the spleen of one individual C57BL/6 mouse were sorted into TCRb+ (T-cell), CD19+ (B-cell) and F4/80+ (macrophage) cell fractions by Flow Cytometry (see Fig. 5A) and then these populations were directly assessed by sc-ICP-MS. Mass-frequency histograms of the background corrected single cell iron data for each cell type are shown in Fig. 5B. T-cells and B-cells had similar geometric mean iron content (Figs. 5B and 5C), corresponding to previous mathematical modelling of lymphocyte cellular iron contents (*29*). The majority of lymphocytes in these populations are non-activated, and their iron content is ∼3-4 fold less than T-cells activated *in vitro* (Fig. 3). This is consistent with the strong upregulation of CD71 that occurs after activation to facilitate iron acquisition, and with mathematical modelling of increased iron content of T-cells in the 1-3 days after activation (*30*, *31*).

**Fig. 5.**
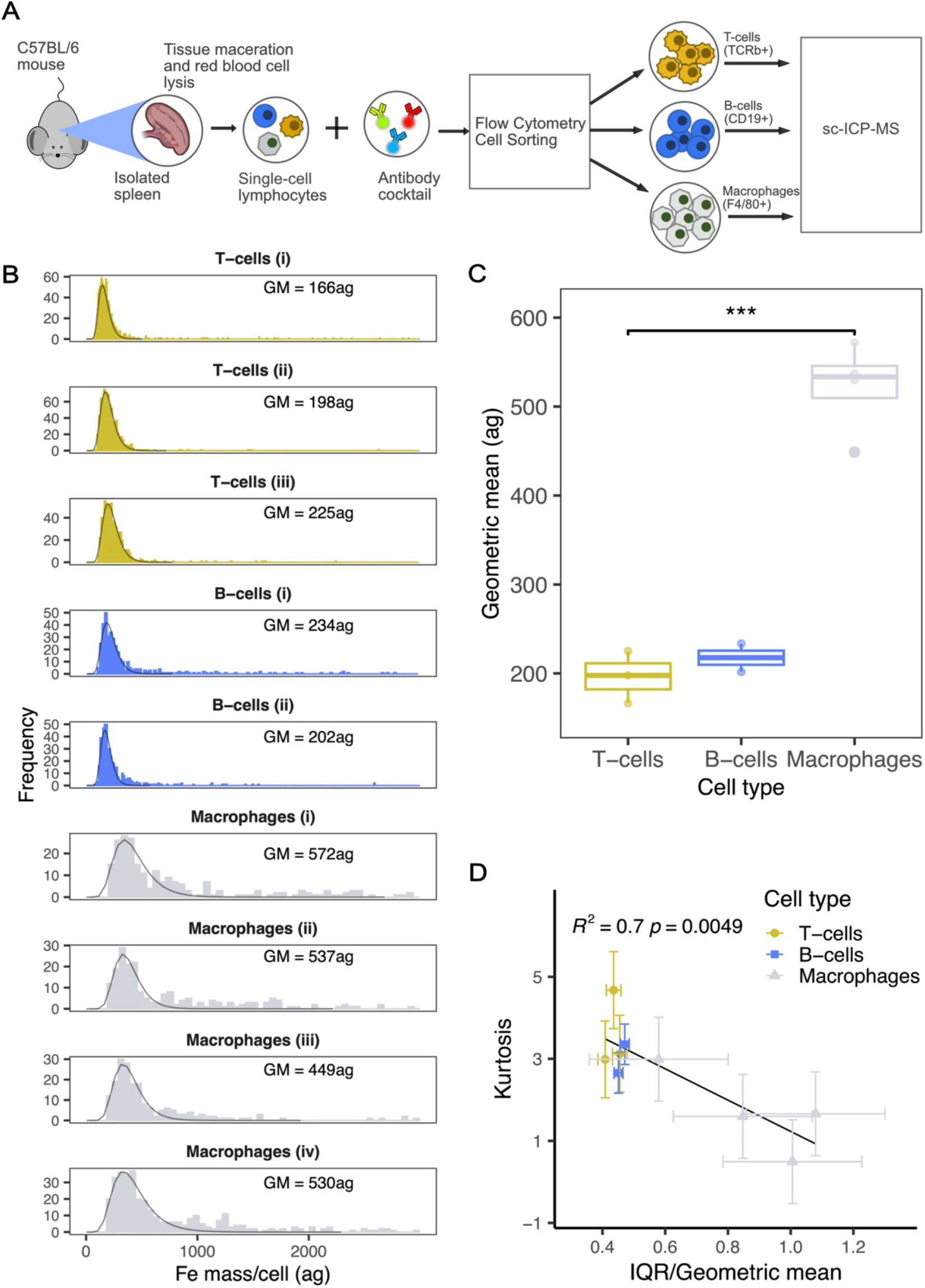
Iron metallomics from unprimed murine splenocytes. **(A).** Experimental design for the assessment of Fe in murine splenocytes that were sorted by Flow Cytometry; **(B).** Histograms showing the mass-frequency distributions of Fe mass/cell for each cell type (and replicate(s)) taken from the mouse, together with lognormal fitting lines. Geometric means (GM) of each population are annotated per facet; **(C).** Boxplot presenting the geometric means from each distribution and for each cell type. Welch’s 2-sample t-test was used to calculate *p-*value for T-cells [n=3] vs Macrophages [n=4]; **(D).** Correlation plot of the kurtosis from each distribution against their interquartile range (IQR)/ geometric mean ratio, together with 1-sigma uncertainty error bars.

Macrophages contained at least twice as much iron per cell (geometric mean) than T-cells or B-cells. Fig 5C shows a boxplot combining each calculated geometric mean showing a statistically significant difference between and T-cells. Macrophages also had broader mass distributions, where replicate measurements attained interquartile ranges extending from 260ag – 572ag; contrasting to equivalent values of 68ag – 98ag and 91 – 110ag, from the T-cell and B-cell populations, respectively (Table S3).

Metallomic heterogeneity in the cell populations from this experiment was evaluated from the kurtosis and ratios of interquartile range/ geometric mean from each distribution (Fig 5D). Kurtosis reports the concentration of data around the mean of a distribution (*32*), where high values suggest a narrow distribution with most data points concentrated around the central mean. Interquartile range/ geometric mean ratios provide a marker of the spread of data in a distribution. The relationship between each cell population and their respective replicates forms a significant inverse correlation (R^2^ = 0.7) (Fig. 5D), with a wide spread in quantitation. T-cells and B-cells collocate in the top left of the correlation plot – suggestive of their narrow distributions and high degrees of peakedness (shown in Fig. 5B), where kurtosis values of around 3 (indicative of a normal distribution), together with contracted interquartile ranges (∼40% of their geometric means) are presented (Fig. 5D). In contrast, macrophages display a much wider distribution that progresses towards values of lower kurtosis and higher IQR/ geometric mean, where 3 out of 4 distributions report kurtosis levels of 1 or below, and 2 out of 4 present distributions that are wider than their presiding geometric means (IQR/ geometric mean >1) (Fig. 5D). This shows the value of sc-ICP-MS to demonstrate variability in heterogeneity of iron content between immune cell types derived from *in vivo* tissue.

### Precise single-cell iron mass distributions from human primary B-cells

To move beyond murine systems, we examined primary B-cells extracted from the peripheral blood of three healthy human donors. After three days of in-vitro culturing in R10 media, iron content of cells was measured by sc-ICP-MS, and flow cytometry was used to assess B-cell purity by analysis of CD19 surface protein expression (see Fig. 6A).

**Fig. 6.**
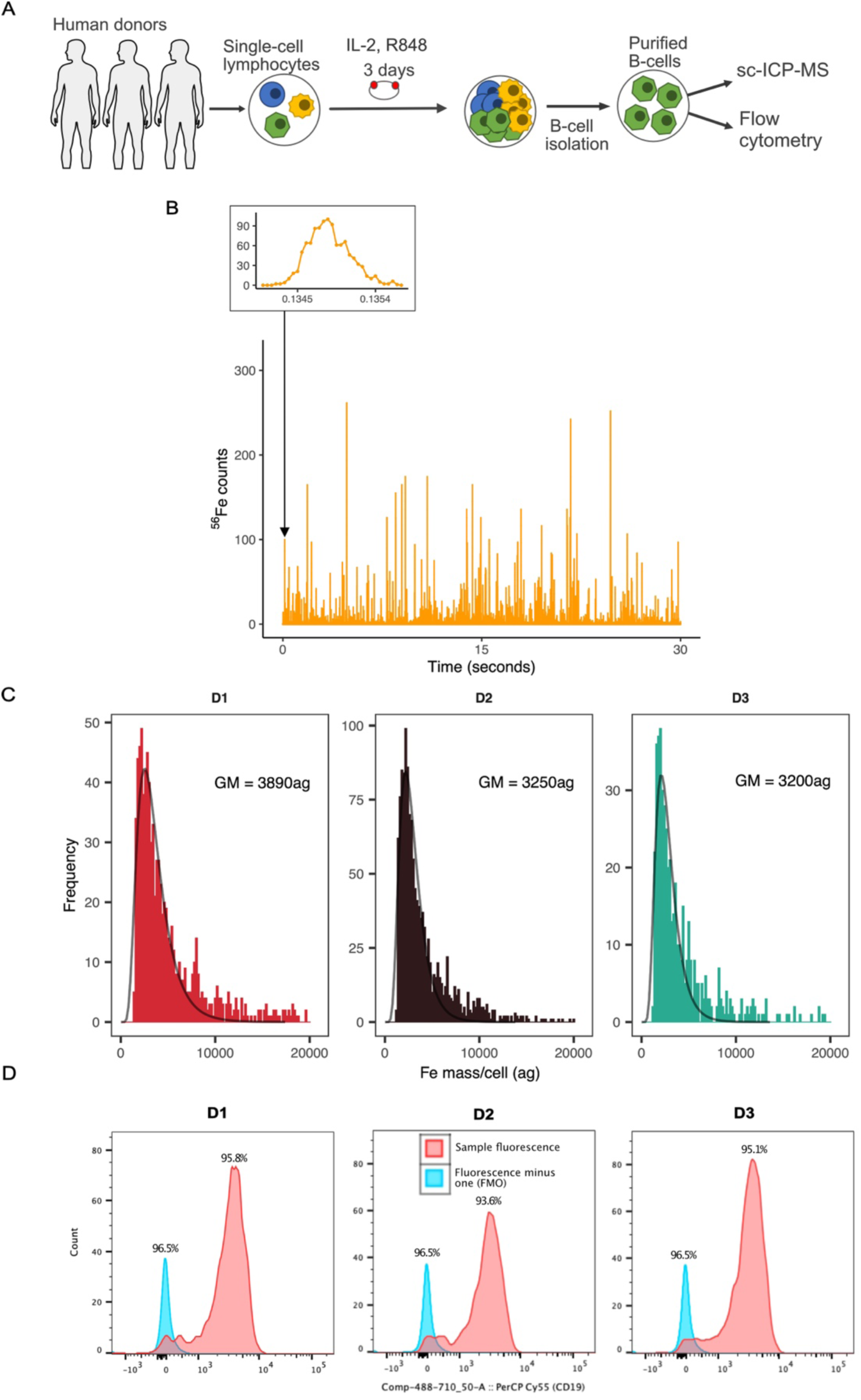
Iron content of primary human B-cells. **(A).** Experimental design for the assessment of Fe in human B-cells. **(B).** An excerpt of time-resolved mass spectra measured for Fe by PerkinElmer NexION 5000 sc-ICP-MS from human B-cells taken from donor D1. Each spike represents a cell metallomic event, where their individual areas correlate to mass of Fe within each cell. The metallomic plumes take ∼0.001s to transit through the instrument (shown in the inset peak). The inset peaks also illustrate the resolutions achieved for the event profiles, where measurements were taken at intervals of 50µseconds. **(C).** Mass frequency histograms presenting distributions of single-cell Fe content in human B-cells determined by sc-ICP-MS for each of three donors, D1-D3, respectively. Mass distributions are shown in attograms (ag) (10^-18^g). Geometric means (GM) of each population are annotated per facet. **(D).** Histograms of CD19+ cell counts, determined by Flow Cytometry, for each of three donors, D1-D3, respectively.

An excerpt of the real-time iron metallomic output is presented in Fig. 6B, where, unlike the murine T-cell experiment, much broader ranges in peak size were obtained. This is likely because human B-cells are larger than murine cells. Similar to the murine T-cell analyses, background corrections were applied to remove the presence of ionic and fragmented cell artefacts. Mass frequency histograms of such corrected data for each donor (D1 – D3) are shown in Fig. 6C, which are presented with optimised bin sizes, lognormal fittings and annotated geometric means for each population. Geometric means ranged from 3200 – 3890ag (Fig. 6C and Table S3), within range of values predicted for human B-cells from mathematical modelling(*29*). Metallomic distributions from the three sample populations present similar mass ranges, where variability of their means (measured by 2-sigma propagation) was 772ag. Additionally, notable levels of heterogeneity (as mentioned above) were also obtained, where interquartile ranges from 2400ag – 3200ag were found (Table S3). Additionally, results from Flow Cytometry confirm the high levels of B-cell purity in each cell population, where CD19+ expression was recorded from 93.6% - 95.8% of the cells measured (see Fig. 6D). These data show that activated human B-cells contain more iron than murine immune cell populations.

## Discussion

Among the range of crucial metallic elements in biology, iron plays a particularly important role due to its ability to shuttle electrons and form bonds with different types of ligands. As a result, iron is utilized in many enzymes and proteins(*1*). However, iron deficiency and iron overload disorders are common worldwide. There is a need to accurately quantify cellular iron content and relate this to changes in cell function, across a range of iron availability.

Sc-ICP-MS can quantify cellular metals across the attogram mass range (10^-18^g)(*33*). Additionally, unlike other solution-based analyses, dilutions applied to reduce the cell concentration do not impact on the signal intensity of the cell events, and may help to reduce the dissolved baseline – thus improving signal: noise ratios. In attempting to improve counting statistics and reduced data volumes, some sc-ICP-MS studies collate metallomics from dwell times exceeding the transit time of a cell event (circa. 200-400 µseconds(*34*)). In contrast, we advocate higher scanning frequencies at intervals faster than cell events, a capability dependent on the ICP-MS instrument being used (*35–37*). Such intervals reduce backgrounds, decrease probability of recording doublets and help distinguish “real” cell events from cell fragments/ debris, improving accuracy in metallomic determinations.

Sc-ICP-MS has limitations whereby components of the sample introduction system cause cell losses during analysis. However, by using a dedicated single-cell microflow injection system, we could consistently achieve transport efficiencies of nearly 15% (Table S2). Sc-ICP-MS may not be a requirement for all fields of metallomic research, where bulk approaches may suffice (e.g. (*38*, *39*)). Nevertheless, our intercalation experiments showed inaccuracies from cell counting and lower detection can decrease accuracy when adopting bulk approaches (see Fig. 1F). A further limitation for bulk analysis is the lack of determination of ‘real’ cell results against fragmented entities.

*In vitro* activated murine T-cells cultured across a 625-fold titration of available extracellular iron maintained an overall mean iron content per cell within 20%. This is consistent with well-described molecular mechanisms of iron homeostasis (such as the IRE-IRP system) (*27*) that prevent cellular iron deficiency and potentially toxic iron over-accumulation. Nevertheless, changes within the 20% window corresponded to CD71 surface expression (which is regulated by intracellular iron) and CD25 (which is a marker of cell activation), and in low iron conditions the cells proliferated markedly less. Our previous work has shown that in the same low iron conditions used here, CD8+ T-cell metabolism is drastically altered, with mitochondrial ATP production and membrane potential reduced and TCA cycle progression imbalanced (Teh et al, *Nat Commun* Accepted). Therefore, within the 20% difference of cellular iron observed lies a substantial variation in cellular activity and function; understanding how relatively small changes in cellular iron result in such effects should be investigated. Our proliferation data also suggest, at least for activated T-cells, that a threshold of cellular iron content may exist below which cells do not divide; the nature of this threshold should be determined by future work. The importance of cellular acquisition for T-cells is demonstrated by human genetics, in which a defect that impairs CD71-mediated iron uptake causes immunodeficiency (*40*); how this defect, and others in different genes that also affect iron handling and cause disease, alter iron content of different cell types remains to be explored, as do effects of nutritional iron deficiency on cellular iron.

Lymphocytes stimulated to proliferate increase iron uptake and increase cell size. We found that compared to activated T-cells, T-cells (and B-cells) derived *ex vivo* from murine spleens without activation contained about 4-fold less iron per cell, in line with predictions based on proteomic surveys of T-cells and known numbers of iron-binding proteins (*31*). Splenic macrophages contained more iron, likely due to their role in the red pulp of the spleen of capturing senescent erythrocytes and recycling haem iron. *In vitro* activated human B-cells contained substantially more iron than murine cells types measured, likely because human lymphocytes are larger than their mouse counterparts. Therefore, it is possible that different cell types have distinct ‘set-points’ for their ideal iron content and that this varies depending on activation state, and species. Other sc-ICP-MS studies have looked at iron uptake into bacteria, for example magnetotactic bacteria increased cellular iron content in proportion to increased availability of environmental iron sources (*14*). Additionally, sc-ICP-MS quantified cis-platin uptake in individual cells from a variety of lymphocyte cell-lines (*12*); including cisplatin resistant (A2780cis) and sensitive (A2780) cells (*41*).

Overall, we show that sc-ICP-MS can rapidly and precisely scan the profiles of cell populations can reveal metal content *ex vivo* and *in vitro* across different experimental conditions and with relevance to activation state and proliferation. Although we have focussed on iron and immune cells, the technique can be applied to other metals and other cell types, mammalian or otherwise. By closer coupling of metallomic measurements to functional parameters we can begin to close the gap between element quantification and cellular activity at single-cell level to better understand how metals influence biology.

## Acknowledgments

The authors would thank Elemental Scientific Inc. for the technical support in commencing analysis with the single cell sample introduction system, and the staff of the Department of Biomedical Services, University of Oxford for animal husbandry and welfare. MT, DC, GP and HD. were supported by a U.K Medical research council, MRC Human Immunology Unit grant awarded to HD. (MCU_12010/10). MT was funded by the Clarendon Fund and the Corpus Christi College A. E. Haigh graduate scholarship. PH is supported by the Human Iron Research in Oxford (HIRO) and PerkinElmer.

## Author contributions

Conceptualization: HD, JW, MT, PH

Methodology: PH, DP, MT, MM, GP

Investigation: PH, MT, MM, NS, MZ, DC, HWC, GP

Visualization: PH, GP

Supervision: JW, HD, DP

Writing—original draft: PH, JW

Writing—review & editing: HD, PH, RH, DP, GP, MT, MM

## Data and materials availability

Tabulated data accompanying the intercalation and murine lymphocyte experiments are included in the supplementary data at the end of this manuscript. Please contact the corresponding author for further information about the data presented in this manuscript.

## Materials and Methods

### Jurkat cell line, rhodium and iridium intercalation

Clone E6-1 Jurkats were purchased from ATCC, a clone of the Jurkat-FHCRC cell line (derivative of the original Jurkat cell line). They were cultured in R10 media (RPMI 1640 supplemented with 10% foetal bovine serum, 1% penicillin-streptomycin and 1% glutamine) and incubated at 37°C and 5% CO_2_ in T75 flasks. For the metal intercalation experiments, freshly passaged cells were divided into subsets (each containing cell concentrations of circa. 4×10^6^ cells/mL) and rinsed by centrifugation with Standard Biotools Maxpar^®^ phosphate-buffered saline (PBS), prior to subsequent fixation in 4% paraformaldehyde for 10 minutes at room temperature. Titrated concentrations of either rhodium or iridium intercalators (Standard Biotools’ 500µM rhodium Cell-ID^TM^ or 125µM iridium Cell-ID^TM^, respectively) were then doped into each sample to form intercalation concentration ranges of 0.05 µM – 0.5 µM and 0.005µM – 0.05 µM, respectively. The cell samples were then stored overnight at 4°C to ensure complete penetration of the Cell-ID^TM^ organo-metallic compounds by passive diffusion through the permeated membranes of each cell. The following day, each sample was divided into two, to provide aliquots for both sc-ICP-MS and CyTOF analysis. Prior to analysis, the cells were rinsed with Standard Biotools Maxpar^®^ cell staining buffer by centrifugation to remove any excess metal accumulation from the cell surfaces.

### Mice and T-cell isolation from peripheral blood, iron titration and *in-vitro* proliferation

OT-I mice (2 x 12-week-old males, and 1 x 13-week-old male), were originally obtained from Audrey Gerard, University of Oxford, and were housed in individually ventilated cages. All animal work was completed under the authority of the UK Home Office project and personal licenses under the Animals (Scientific Procedures) Act (ASPA) 1986. Mice were sacrificed via rising concentration of CO_2_ followed by cervical dislocation. Plates for CD8+ T-cell culture were pre-treated with 5 ug/mL α-CD3 (Biolegend, 100239) in phosphate buffered saline for 2 hours at 37°C. Spleen and lymph nodes were collected from euthanised mice and macerated through 40 μm filters using PBS supplemented with 2% fetal bovine serum and 1 mM EDTA (Invitrogen, AM9260G). CD8+ T-cells were isolated from the single-cell suspension using the EasySep Mouse CD8+ T-cell isolation kit (Stem Cell Technologies, 19853) and the EasyEights EasySep magnet (Stem Cell Technologies, 18103). Isolated cells were stained with cell trace violet (CTV, Invitrogen, C34557) for 8 minutes at 37°C in PBS and then washed. CD8+ T-cells were plated at a concentration of 0.5×10^6^ cells/mL on the α-CD3 pre-treated plates. Cells were grown in iron-free media (RPMI1640 (Gibco, 21875034), 10% iron free serum substitute (Pan Biotech, P04-95080), 1% glutamine (Sigma Aldrich, G7513-100ML) and 1% penicillin/streptomycin (Sigma Aldrich, P0781-100ML)) supplemented with set concentrations of holo and apotransferrin. Human holotransferrin (R&D systems, 2914-HT-001G) was added at concentrations of 0.001 mg/mL to 0.625 mg/mL. Total transferrin levels were kept at a constant concentration of 1.2 mg/mL by adding the appropriate amount of human apotransferrin (R&D systems, 3188-AT-001G). Cells were also treated with 50 μM β-mercaptoethanol (BME, Gibco, 31350-010), 1 μg/mL α-CD28 (Biolegend, 102115) and 50 U/mL IL-2 (Biolegend, 575402) to activate the cells. CD8+ T-cells were cultured at 37°C, 5% CO_2_ for 48h.

After incubation the cells were harvested, counted and aliquots divided between sc-ICP-MS and Flow Cytometry; where ∼ 2×10^6^ cells/mL were retained for sc-ICP-MS. The sc-ICP-MS cell aliquots were then rinsed twice by centrifugation with Standard Biotools Maxpar^®^ PBS, followed by resuspension in 1mL 4% paraformaldehyde for fixation at room temperature for 10 minutes. The samples were then rinsed twice by centrifugation with Standard Biotools Maxpar^®^ Cell Staining Buffer to remove any excess iron remaining from the cell surfaces, followed by resuspension in Standard Biotools Maxpar^®^ Fix and Perm reagent for overnight storage at 4°C.

### Mice, *in-vivo* splenic lymphocyte isolation and cell sorting

Mice were bred and maintained in the University of Oxford’s specific pathogen-free (SPF) animal facilities. Experiments were conducted in accordance with local animal care committees (UK Scientific Procedures Act of 1986) under the project licence PP1487090. Mice were kept in individually ventilated cages with environmental enrichment with a 12h light/dark cycle. The C57BL/6 mice were purchased from Invigo Ltd. and were culled by rising CO_2_ inhalation.

The spleen was isolated from surrounding tissues with tweezers and maintained on ice in PBS/BSA. Tissues were processed to a single cell suspension by maceration through a 70uM mesh filter. Spleen samples were then incubated with 1mL ACK lysis buffer for 3 minutes to lyse red blood cells.

Single-cell suspensions were stained with LIVE/DEAD Fixable Near IR Dead Cell dye and TruStain FcX (anti-mouse CD16/32) antibody, diluted 1:1000 and 1:100, respectively in PBS for 10 minutes on ice in the dark. Cells were then stained in PBS with antibodies purchased from Biolegend for 20 minutes on ice in the dark. Cells were then washed (2 minutes at 1200rpm and 4°C) and fixed in Fixation Buffer for 20 minutes at room temperature in the dark. Samples were then washed and resuspended in Standard Biotools’ Maxpar^®^ Cell Acquisition Solution Plus (CAS+) buffer. Samples were acquired and sorted on a BD FacsAria Fusion using FACs Diva Software.

### Human blood donations: B-cell isolation, purification and *in-vitro* proliferation

Blood samples taken from three healthy donors from the John Radcliffe Hospital, Oxford, United Kingdom, were utilised for single-cell iron analysis in B-cells by sc-ICP-MS. Each sample was collected after obtaining written consent and ethical approval from the University of Oxford’s Central University Research Ethics Committee (CUREC). The samples were collected in EDTA, which was followed by PBMC isolation by density gradient centrifugation: Greiner Bio-One Leucosep tubes, containing 15mL of Lymphoprep (Stem Cell Technologies) and collected blood were centrifuged at 1000 x g for 1 minute at ambient temperature. EDTA blood was extracted into the upper chamber of the Leucosep tube and centrifuged at 1000 x g for 15 minutes with no brake. The cloudy buffy-coat layer, containing PBMCs, was extracted, and the cells were subsequently rinsed twice with R0 media (RPMI 1640 supplemented with 1% penicillin-streptomycin and 1% glutamine) and PBS, respectively. Cell Trace Violet (Thermo Fisher Scientific) was added to the rinsed PBMCs as a tracer for proliferation and incubated at 37°C in 5% CO_2_ for 8 minutes. After incubation, the cells were rinsed with R10 media, counted and then diluted to 8×10^6^ cells per mL in R10 media. Two million cells were subsequently added per well into a 24-well rounded bottom plate, together with aliquots of 0.25mL of R10 media and 0.5mL of R10 media supplemented with 1µg/mL R848 (Stem Cell Technologies) and 10 ng/mL of recombinant IL-2 (PeproTech). The prepared cells were then cultured for 3 days at 37°C in 5% CO_2_. Following polyclonal stimulation, the cells were harvested, washed in R10 media and counted. B-cells were then purified from the other harvested PBMCs by negative selection using a Human B-cell Isolation Kit (Stem Cell Technologies) according to the manufacturer’s instructions. Following purification, the B-cells were transferred to a 96-well rounded bottom plate and rinsed with PBS, following Fc Receptor (FCR) blocking and live/dead staining. This was followed by the subsequent labelling of the cells with combinations of anti-CD19-PerCP Cy5.5, anti-CD21 (Alexa Fluor 700), anti-CD27 (PE-Cy7), anti-CD38 (BV510), anti-CD69 (BV605), and anti-CD71 (PE/Dazzle 594) in PBS in addition to incubation for 20 minutes on water ice and fixation buffer (Biolegend). Before intracellular staining with anti-IgG (BV711) and anti-IgD (FITC) the cells were permeabilised with perm buffer for 20 minutes on water ice. Prior to measurement by Flow Cytometry, fluorescence minus one controls (FMOs) were included for each marker, in addition to an unstained control.

### Single-cell inductively coupled plasma mass spectrometry (sc-ICP-MS)

Following all of the methodologies described above, the prepared cell suspensions were also rinsed three times in Standard Biotools’ Maxpar^®^ Cell Acquisition Solution Plus (CAS+) prior to sc-ICP-MS analysis. This was undertaken to ensure the removal of any remaining residually-retained metals from the cell surfaces, and to an exchange into a suspension media suitable for analysis by this technique. Moreover, CAS+ is proven to be an optimal choice for analysis utilising our analytical setup over other commonly used carrier reagents(*42*, *43*), where its combination of a neutral pH in addition to a higher ionic content than water provides a higher analytical performance for metallomic analysis when combined with a wider bore injector. Proceeding the CAS+ rinses, the final suspensions were filtered through 35µm nylon mesh filters, and the subsequent cell suspensions counted, diluted to 10^5^ cells/mL cell concentrations (if required) and then immediately measured by sc-ICP-MS. Supernatants from the final rinse cycles of selected samples were retained, to test for any leakage of intracellular metals.

All sc-ICP-MS measurements were conducted using either a NexION 5000 multi-quadrupole ICP-MS (PerkinElmer) or a NexION 350D ICP-MS (PerkinElmer), in time-resolved mode (see instrument conditions and further details in Table 1). Both instruments were equipped with an Elemental Scientific Inc. single cell introduction system, which comprised of a CytoNeb 50 nebuliser, a CytoSpray linear pass spray chamber, a 2.0mm tapered injector (PerkinElmer White Cassette torch with 2.0mm injector for the NexION 5000) and a micro*FAST* autosampler (which provided an additional final agitation of the suspension using its ‘Mix’ submethod to ensure homogenisation of the aliquoted cell suspension for analysis). This apparatus, like many others, also used for single-cell metallomics research(*12*, *22*, *44*), was essential for this analysis to ensure the highest levels of cell transmission to the instrument, where micro-flow volume injections of cells were analysed for precise single-cell metallomics by the mass spectrometer from discrete measurements of their resultant ionic plumes.

The principle of sc-ICP-MS relies on each individual cell from a population to be sequentially introduced to the instrument, where they are individually vapourised, atomised, and their metallomic constituents ionised for measurement within durations of around 200-400 µseconds(*34*). Known as a “cell event”, the instrument must be able to capture the measurement of the metallomic plume for a particular element during its transit through the mass spectrometer. Data output was evaluated using PerkinElmer’s Syngistix Single Cell software module, where the peak area from each cell event was calculated to determine the mass of metal per cell. Further details relating to the optimisation of this instrument setup are described in the supplementary text.

### Bulk Inductively coupled plasma mass spectrometry (bulk-ICP-MS)

Bulk ICP-MS analysis utilised cell aliquots remaining after the metal intercalation sc-ICP-MS analyses, where approximately 0.2×10^6^ cells were dissolved in 2% v/v HNO_3_ within metal-free centrifuge tubes at ambient temperature for 48 hours. Measurements were conducted using the NexION 350D ICP-MS (PerkinElmer) using instrument conditions described in Table 2.

### Mass Cytometry (CyTOF)

Mass Cytometry was used in this study to validate metallomics data obtained by sc-ICP-MS from the metal intercalation experiments on Jurkat cells (as described above). In an analogous approach to the sc-ICP-MS analyses, the prepared cell subsets allocated for Mass Cytometry measurements were rinsed three times by centrifugation with CAS+ to entirely remove any surface-retained metals and to transfer into the instrument carrier media. All measurements using this technique were conducted using a Standard Biotools Helios CyTOF, which employed its standard single cell suspension pneumatic sample introduction system. The mass cytometer was tuned, and its performance confirmed using EQ^TM^ Four Element Calibration Beads. The cells were diluted to 10^6^ cells/mL in CAS+ with 10% EQ^TM^ Four Element Calibration Beads. The .fcs files were acquired and processed (including normalisation) in the CyTOF Software v.7 (Standard Biotools).

### Flow Cytometry

Flow Cytometry analysis was incorporated into this study to validate the iron metallomic data obtained by sc-ICP-MS. Cells were transferred to 96-well round bottom plates and washed with PBS. Cells were stained with an antibody cocktail, and the Zombie NIR fixable viability kit (1:1000, Biolegend, 423105) prepared in PBS for 20 minutes on ice. Cells were subsequently fixed with 2% paraformaldehyde (Pierce, 28906) for 20 minutes on ice, washed and resuspended in PBS. Cells were analysed on either a BD Biosciences LSR Fortessa^TM^ X50 instrument or a Attune NxT flow cytometer (Thermofisher Scientific), and the data evaluated using FlowJo (BD biosciences) software.

### Statistical Analysis

All single-cell metallomics data was evaluated using the single-cell module within PerkinElmer’s Syngistix^TM^ ICP-MS software. Complimentary geometric means were calculated in Excel. Other linear regression determinations, such as linear fitting equations, R^2^ and *p* values were calculated in R, utilising the dplyr and ggplot2 libraries.

## Supplementary Materials

### Supplementary Methods

#### Background correction of sc-ICP-MS data

Single cell inductively coupled plasma mass spectrometry (sc-ICP-MS) provides precise analysis of individual cells from entire cell populations for their intracellular metal contents – a promising and powerful metallomic analytical tool due to its high sensitivity and rapid measurement capabilities. Moreover, the high-resolution time-series data generated by this technique allows the user to extrapolate a wealth of metallomic information, including heterogeneity within populations. Nevertheless, it is essential that this near-constant data-stream is corrected for the measurement baseline, which may comprise either dissolved media, and/or constituent fragmented or below detection cell readings. The dissolved component is recorded as a constant signal that is elevated above baseline (*26*), which if stable, can be systematically deducted by 3-sigma + mean background corrections – as performed by the Single Cell module within the PerkinElmer Syngistix software in this study. However, correcting for concurrent irregular baselines caused by fragmented cells/ cells below detection, involves a more methodical approach, requiring detailed inspections of the peak detail. By conducting sc-ICP-MS at short measurement intervals (<0.1ms), we uncovered highly constrained durations for whole cell events (∼1ms when using NH_3_ cell gas and ∼0.5ms when using no cell gas); contrasting to events with distinctly shorter time intervals near to the baseline (fragments or cells with extremely low metal content). Therefore, by virtue of the recorded event duration times, we were able to discriminate real whole cell metallomic events from cell fragment artifacts/ cells below detection. In the experiments that we conducted in this study, our threshold was selected by the lowest peak amplitude that was able to sustain a whole cell event. Examples of both peak types are presented in Fig. S1, where the highlighted “non-cell” event exhibits a duration time that is > 30% narrower than the example “cell event”.

Verification of the accuracy in our background correction methodology is highlighted by the distinct similarities in the transport efficiency data obtained from the murine T-cell experiment, where average calcium and iron-derived values were 12.2% and 12.7%, respectively (see Table S2). Furthermore, these values also replicate transport efficiencies reported from similar studies (*12, 13, 41*), thus further validating our background correction method.

### Supplementary Figures Figs. S1 – S3

**Fig. S1.**
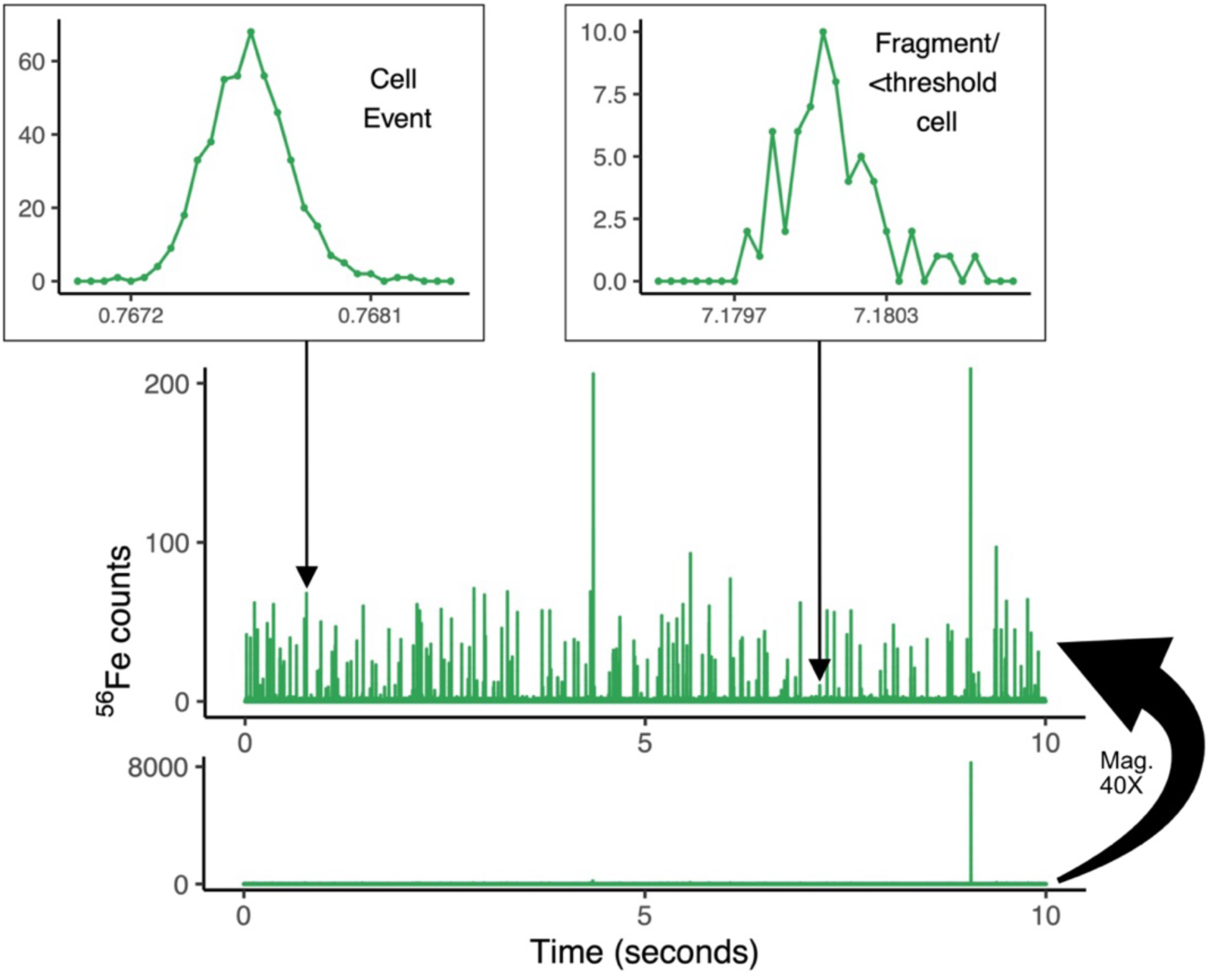
Time-resolved single cell mass spectra measured by SC-ICP-MS from murine T-cells. Measurement of iron in T-cells taken from mouse 3 and the 0.005mg/mL condition. Each spike represents an individual metallomic event, resolved as a peak (see cell event inset), where areas under each peak correlate to intracellular iron content. The left inset, taken at ∼0.76 seconds highlights a whole cell event with a peak duration of ∼1ms. The right inset, taken at ∼7.18 seconds denotes a fragment from a lysed cell/ cell event below detection with a distinctly shorter peak duration of (∼0.65ms).

**Fig. S2.**
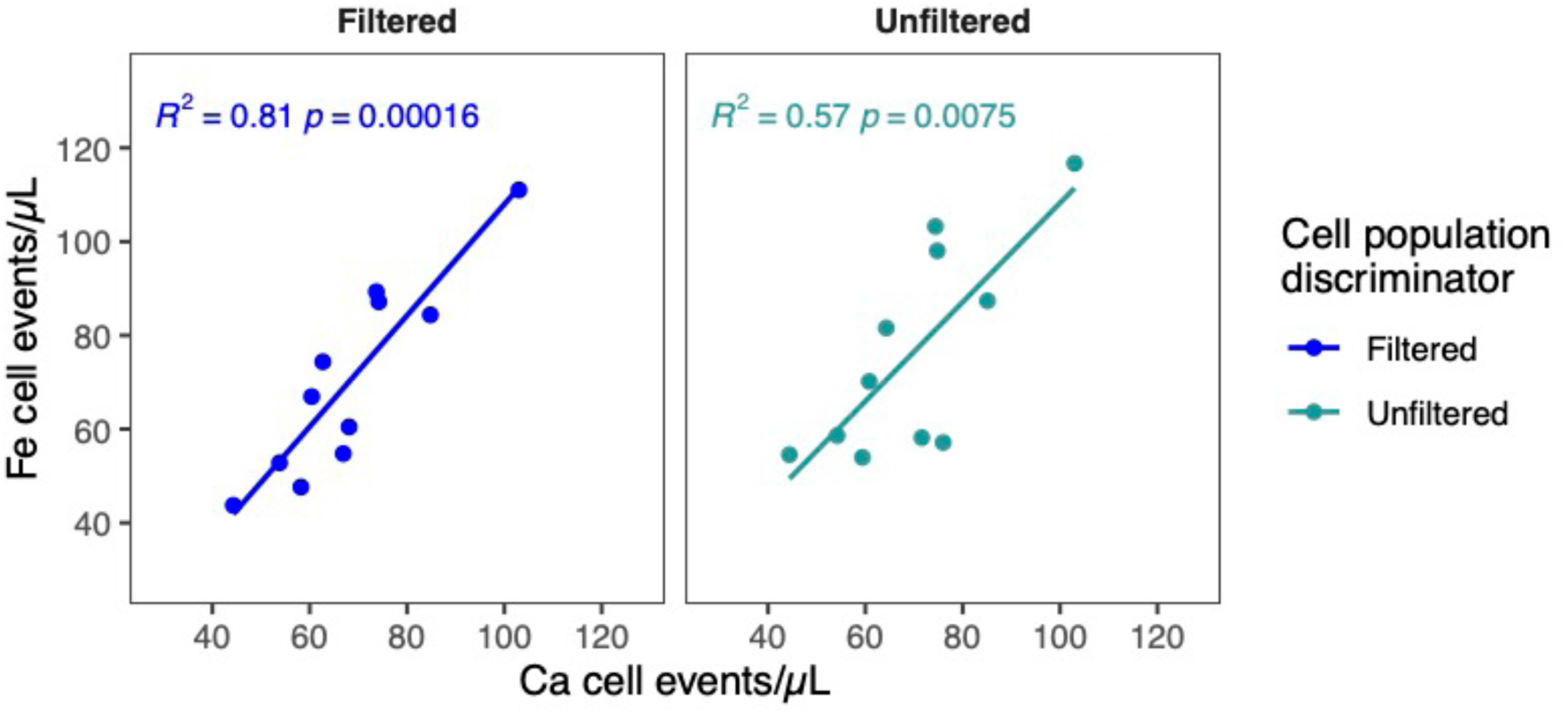
Correlation of measured cell event frequencies (iron vs calcium) for both filtered (events constrained within lognormal distributions) and unfiltered (all recorded data above background threshold). Values displayed are the number of events calculated per µL of sample measured by the sc-ICP-MS. All data are exclusive of outliers (calculated as >2-sigma from a series of measurement replicates [n=4]) and includes linear trendlines, R^2^ and *p*-values.

**Fig. S3.**
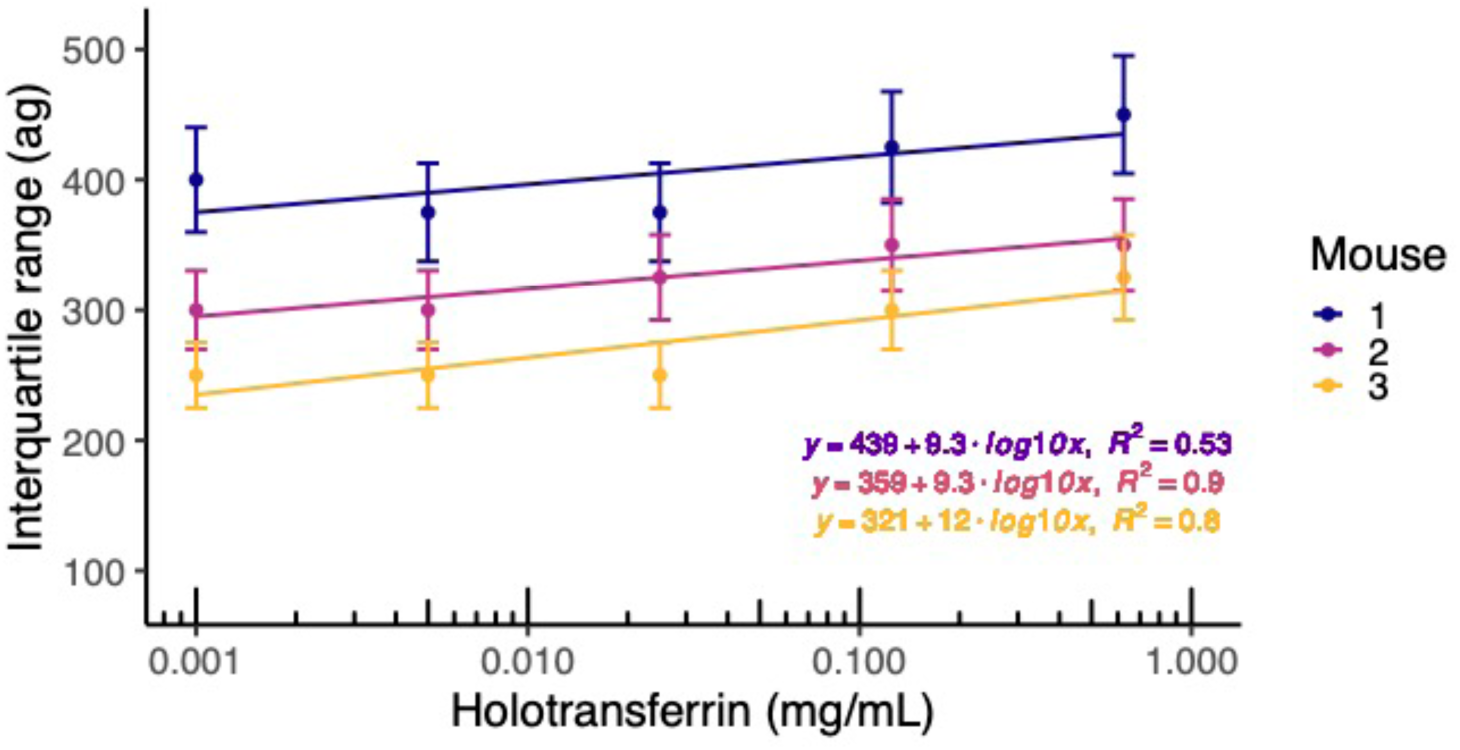
Correlation of inter-quartile ranges from each sample distribution against holotransferrin iron-media condition in the murine T-cell experiment. Trend lines, associated equations and R^2^ values displayed to show logarithmic increases with iron condition. Error bars present 2-sigma uncertainty calculated from replicate measurements [n=4].

### Supplementary Tables S1 – S4

**Table S1.** Statistical evaluation of SC-ICP-MS data from the metal-intercalation experiments. All above data calculated from extrapolated metal-intercalated metallomic distributions. Cells were measured for 1 minute time periods. *Measured at a two-times higher dilution.

| Intercalation molarity (μmol) | Metal intercalator | Number of cell events | Geometric mean (ag) | Interquartile range (ag) |
| --- | --- | --- | --- | --- |
| 0.005* | Iridium | 15 | 44 | 21 |
| 0.01 |  | 24 | 26 | 7.5 |
| 0.02 |  | 184 | 28 | 15 |
| 0.03 |  | 382 | 44 | 24 |
| 0.04 |  | 479 | 54 | 27 |
| 0.05 |  | 577 | 64 | 36 |
| 0.05 | Rhodium | 15 | 129 | 50 |
| 0.1 |  | 63 | 158 | 48 |
| 0.2 |  | 268 | 219 | 104 |
| 0.3 |  | 774 | 502 | 225 |
| 0.4 |  | 424 | 580 | 320 |
| 0.5 |  | 442 | 760 | 385 |

**Table S2.** Transport efficiency data from the murine T-cell experiments. All transport efficiency data calculated by the particle frequency method (*26*). Both iron and calcium-derived transport efficiencies were derived directly from the murine T-cells, and holmium-derived transport efficiencies from EQ^TM^ Four-Element Calibration Beads.

| Mouse | Holotransferrin condition (mg/mL) | Transport efficiency: T-cell Fe-derived (%) | Transport efficiency: T-cell Ca-derived (%) | Transport efficiency: EQ™ Four-Element Calibration Beads: Ho-derived (%) |
| --- | --- | --- | --- | --- |
| n/a | n/a | -/- | -/- | 29.1 |
| n/a | n/a | -/- | -/- | 22.9 |
| 1 | 0.001 | 9.6 | 10.8 | -/- |
|  | 0.005 | 7.8 | 11.8 | -/- |
|  | 0.025 | 10.3 | 8.8 | -/- |
|  | 0.125 | 16.3 | 8.8 | -/- |
|  | 0.625 | 19.0 | 15.7 | -/- |
| 2 | 0.001 | 8.1 | 13.2 | -/- |
|  | 0.005 | 21.6 | 18.2 | -/- |
|  | 0.025 | 10.4 | 10.6 | -/- |
|  | 0.125 | 17.0 | 17.2 | -/- |
|  | 0.625 | 12.6 | 15.5 | -/- |
| n/a | n/a | -/- | -/- | 25.9 |
| 3 | 0.001 | 10.8 | 10.9 | -/- |
|  | 0.005 | 11.1 | 7.3 | -/- |
|  | 0.025 | 15.7 | 14.5 | -/- |
|  | 0.125 | 7.9 | 9.7 | -/- |
|  | 0.625 | 11.5 | 10.4 | -/- |
| n/a | n/a | -/- | -/- | 28.9 |
|  | **Average** | **12.7** | **12.2** | **26.7** |

**Table S3.**
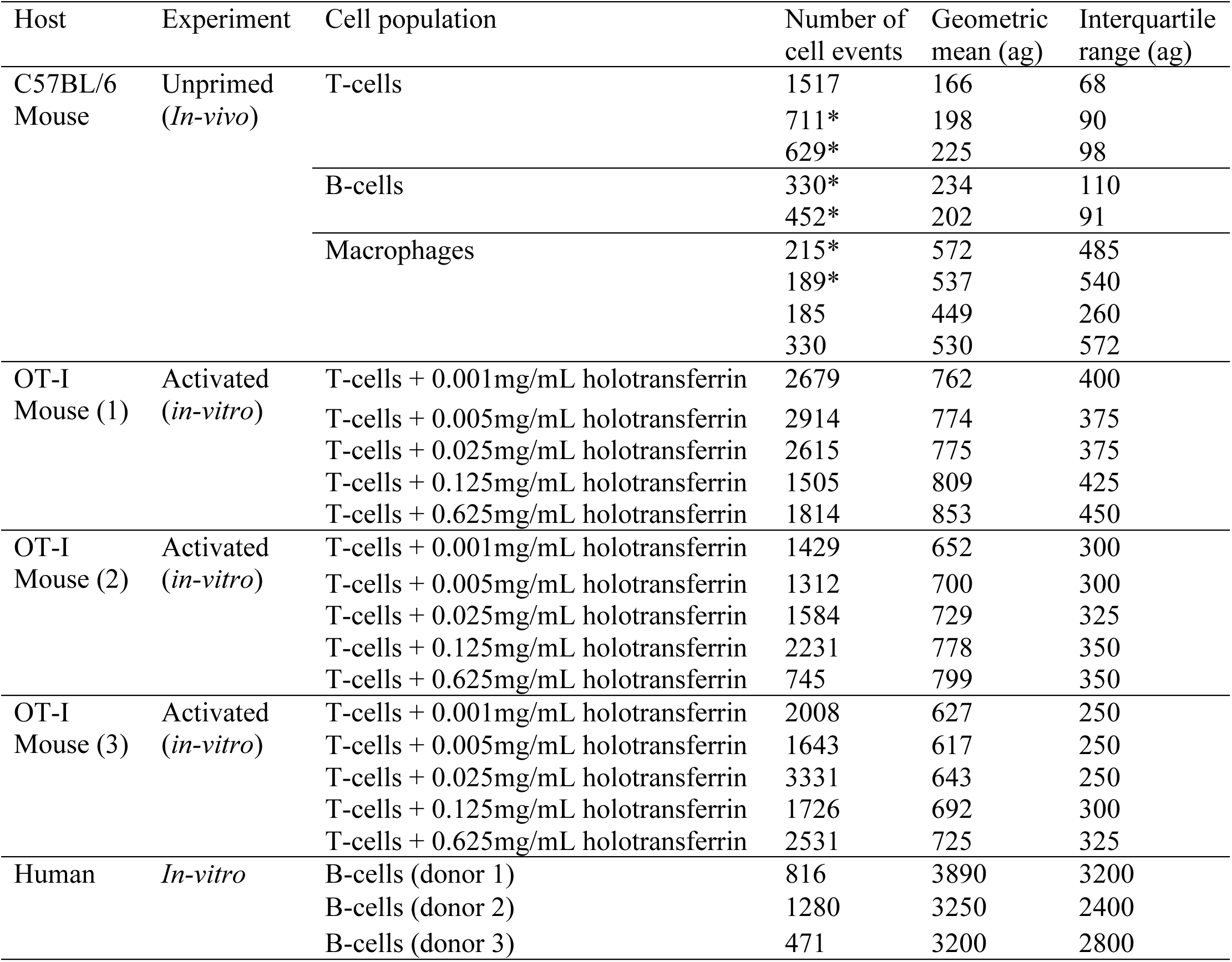
Statistical evaluation of SC-ICP-MS data from iron metallomics experiments. All above data calculated from extrapolated iron metallomic distributions. Cells were measured for 3-minute time periods for the activated T-cell experiments and human B-cells; and 4 minutes for the unprimed cells. *measured at a two-times dilution.

**Table S4.** Live cell counts and total number of cells harvested following the 2-day culturing period for the murine T-cell experiment.

| Mouse | Holotransferrin<br>condition<br>(mg/mL) | Live cell<br>count A<br>(cells/mL) | Live cell<br>count B<br>(cells/mL) | Mean (A<br>and B)<br>(cells/mL) | Total<br>volume<br>(mL) | Total<br>number of<br>cells |
| --- | --- | --- | --- | --- | --- | --- |
| 1 | 0.001 | 1.99E+05 | 2.77E+05 | 2.38E+05 | 7.5 | 1.79E+06 |
|  | 0.005 | 2.67E+05 | 3.40E+05 | 3.04E+05 | 7.5 | 2.28E+06 |
|  | 0.025 | 3.56E+05 | 2.72E+05 | 3.14E+05 | 7.5 | 2.36E+06 |
|  | 0.125 | 2.04E+05 | 2.25E+05 | 2.15E+05 | 7.5 | 1.61E+06 |
|  | 0.625 | 2.30E+05 | 2.83E+05 | 2.57E+05 | 7.5 | 1.92E+06 |
| 2 | 0.001 | 1.88E+05 | 2.77E+05 | 2.33E+05 | 6 | 1.40E+06 |
|  | 0.005 | 3.61E+05 | 3.98E+05 | 3.80E+05 | 6 | 2.28E+06 |
|  | 0.025 | 3.35E+05 | 3.03E+05 | 3.19E+05 | 6 | 1.91E+06 |
|  | 0.125 | 2.20E+05 | 2.67E+05 | 2.44E+05 | 6 | 1.46E+06 |
|  | 0.625 | 2.93E+05 | 2.56E+05 | 2.75E+05 | 7.5 | 2.06E+06 |
| 3 | 0.001 | 4.92E+05 | 5.28E+05 | 5.10E+05 | 7.5 | 3.83E+06 |
|  | 0.005 | 5.91E+05 | 5.02E+05 | 5.47E+05 | 7.5 | 4.10E+06 |
|  | 0.025 | 4.39E+05 | 5.81E+05 | 5.10E+05 | 7.5 | 3.83E+06 |
|  | 0.125 | 4.81E+05 | 4.50E+05 | 4.66E+05 | 7.5 | 3.49E+06 |
|  | 0.625 | 3.92E+05 | 4.08E+05 | 4.00E+05 | 7.5 | 3.00E+06 |

